# Improving the Safety of *N,N*-Dimethylacetamide (DMA) as a Potential Treatment for Preterm Birth in a Pregnant Mouse Model Using a Vaginal Nanoformulation

**DOI:** 10.1101/2025.01.16.633348

**Authors:** Asad Mir, Teeshavi Acosta, Marta Concheiro-Guisan, Steven M. Yellon, Ketan Patel, Sandra E. Reznik

**Author notes:** **Corresponding Author:** Sandra E. Reznik, MD, Ph.D; Department of Pharmaceutical Sciences, St. John’s University, Queens, NY, 11439, USA.

## Abstract

Vaginal administration and the uterine first pass effect allow for preferential delivery of drugs to the reproductive tract. Dimethylacetamide has previously been shown to delay preterm birth in a pregnant mouse model when given intraperitoneally but the effectiveness of a vaginal nanoformulation of dimethylacetamide has yet to be tested. The purpose of this study was to compare the two formulations of dimethylacetamide for efficacy in rescuing pups from preterm birth in an inflammation-induced mouse model, effects on the maternal fetal interface, and pharmacokinetic profiles in maternal plasma. Timed pregnant CD1 mice were given a 1.56 mg/kg intraperitoneal dose of lipopolysaccharide followed by 3 doses of either vaginal dimethylacetamide or intraperitoneal dimethylacetamide. Mice were monitored for 48 hours and times of deliveries were recorded. Additionally, CD1 mice in late gestation were given a single dose of either vaginal or intraperitoneal dimethylacetamide and blood was drawn at 3 different time points following administration. Vaginal administration of dimethylacetamide had similar efficacy in delaying inflammation induced preterm birth as intraperitoneal administration but resulted in lower concentrations in the systemic circulation and decreased effects on the maternal fetal interface. Vaginal nanoformulations should be explored for their potential therapeutic value for the delay of preterm birth.

## Introduction

Preterm birth (PTB) continues to be one of the leading causes of neonatal morbidity and mortality worldwide^1–3^. In the post-approval confirmatory trial for Makena®, there was no compelling evidence for the drug to reduce PTB rates^4^ and it was withdrawn in 2023^5^. Recent studies suggest that utilizing the vaginal route of administration shows promise^6–9^ in that drugs administered this way undergo the “first uterine pass effect”, an aggregation of biochemical and physiological mechanisms working together to deliver therapeutic agents to the uterine-cervical-vaginal region at higher concentrations when compared to other routes of administration^7,10^. Although this phenomenon has been well-documented for years^6,11^, it wasn’t until recently that vaginal administration was being touted as a viable alternative to traditional routes of administration^12–16^. This is due to the physical and chemical properties of many small-molecule therapeutics that make it unlikely for them to traverse the physical and immunological barrier present at the site of the vaginal opening, specifically the protective mucus layer secreted by the cervix^17–19^. However, in the last decade, studies have demonstrated that this obstacle can be circumvented using nanomedicine^9,12,20^. Nanoformulations can be optimized so that they can penetrate the mucus barrier of the vagina and unload their cargo into the reproductive tract^21–23^. Many small molecule lipophilic candidates whose physiochemical properties previously impaired their solubility, pharmacokinetics and overall efficacy can be revisited now that they can be loaded into a nanoformulation^24–26^.

Whether at term or preterm, the process of parturition is analogous to multiple inflammatory processes superimposed over one another^27,28^, with physiological changes such as cervical ripening and uterine contractions signaling the arrival of the conceptus^29,30^. Previous studies have shown that the common pharmaceutical excipient, *N,N-*dimethylacetamide (DMA) can delay and prevent preterm birth in an inflammation-induced mouse model via attenuation of the NF-κB-pathway^31,32^. In our study, DMA was loaded into a self-nanoemulsifying drug delivery system (SNEDDS) and given to pregnant CD-1 mice with its therapeutic and adverse effect(s) compared to the traditional formulation given intraperitoneally.

For treating ailments in the reproductive tract, it is believed that vaginal administration is not only a viable option but a favorable alternative to other more traditional routes of administration. Therefore, the main aims of this study were: 1) to assess the efficacy of two different formulations of DMA to rescue developing pups from preterm birth and spontaneous abortion in an inflammation-induced CD1 mouse model; 2) to classify adverse and toxic effects in select tissues of the female mouse reproductive tract; 3) to compare levels of DMA in the systemic circulation at different time points following administration.

## Materials & Methods

### Animals

The study protocol and all procedures complied with the St. John’s University College of Pharmacy and Health Sciences Animal Care and Use Committee. Research was conducted in accordance with the NIH Guide for the Care and Use of Laboratory Animals, eighth edition, National Research Council (US) Committee for the Update of the Guide for the Care and Use of Laboratory Animals. Eight-week-old female CD-1 mice were purchased from Charles River (Wilmington, MA) and housed in the university’s animal care center with 12-hour light/dark cycles, being given food and water *ad libitum*. For mating, 2-3 female dams were placed in a cage with one male mouse overnight. Successful mating was confirmed by detecting vaginal plugs in the female dams and pregnancy was confirmed one week later by examining the abdomen and measuring the weight of the female mice.

### Efficacy study and comparative study

For assessing the efficacy of both formulations to prevent or delay preterm birth in an inflammation-induced mouse model, pregnant mice were given an intraperitoneal (ip) injection of lipopolysaccharide (LPS) (Sigma-Aldrich, St. Louis, MO; Serotype O55:B5,) at a dose of 1.56 mg/kg on gestational day 15.5 (E15.5) at a time designated as time zero (T0). Mice were given ip DMA (ThermoFisher Scientific, Fair Lawn, NJ) at 1.6 g/kg at the following time points; T minus 15 minutes, T plus 8 hours and T plus 12 hours and monitored for 24 hours to record time of delivery and time of pups dropping. Mice given 0.94g/kg of SNEDDS DMA (St. John’s University, New York, NY) were dosed at T minus 15 minutes, T plus 8 hours and T plus 32 hours and monitored for 48 hours to record time of delivery and time of pups dropping. For the comparative study, the above procedure was repeated but without LPS to ascertain adverse effects in the reproductive tract.

### Histology/immunohistochemical staining and semi-quantitative analysis

H&E staining and immunohistochemical staining of the placenta and yolk sac were performed as previously described^33,34^. ImageJ/Fiji software version 2.14.0 (National Institutes of Health) was used to analyze digitized images (400x magnification, 438.2 μM x 438.2 μM) of H&E-stained and CD31-stained sections of the placenta. Semi-quantitative analysis was based on square area of colored pixels (blood/red, CD31/brown). The “*color deconvolution”* plugin was used on the digitized images to unmix the dyes in the RGB image using the “H&E” vector and separating the image into its individual red, blue and green channels via an NIH macro kindly provided by A.C. Ruifrok. The red channel was used to subtract out the background and measure the square area taken up by red blood cells. For quantifying CD31-immunostained sections, color deconvolution was done using the “*H DAB”* vector to unmix the dyes and separate micrographs into individual brown, blue, and green channels with the brown channel being used to subtract out the background and measure the square area of CD31 localization in the labyrinth. Semi-quantitative analysis workflow is depicted in supplemental figure 2.

### Placental genotyping, sex determination and sex as a biological variable

Placenta samples were deparaffinized with xylene and rehydrated with a series of ethanol washes as previously outlined^35^. Tissue section(s) 3 mm in diameter were placed into a Transnetyx® wellplate and shipped to Transnetyx®. DNA isolated from the placental tissue was treated with their gender-determining “Y” probe to determine the sex of the developing pup attached to the placenta.

### Plasma collection and LC/MS-MS analysis

E15 mice were given either ip DMA or vaginal DMA at a dose of 1.56 g/kg. Blood was obtained at three different time points by puncturing the submandibular vein with a 4 mm Goldenrod™ lancet (Medipoint, Mineola, NY) and holding the mouse over a heparin-coated microvette as blood dripped from the cheek. Microvettes were then placed in a 4°C centrifuge for 20 minutes at 2000 rcf with the supernatant being collected and stored at -80°C. A previously validated method for quantifying DMA in pediatric plasma was adapted to analyze DMA levels in mouse plasma^37^.

Separation of DMA and internal standard was achieved using a Shimadzu LC30-AD system. Elution of analytes was performed using a ThermoFisher Scientific HyperCarb (100 mm x 2.1 mm x 3 μM) stationary phase. The mobile phase consisted of 0.1% formic acid in water (solvent A) and methanol (solvent b). Separation was achieved using a gradient method starting at 5% solvent b and finishing at 95% solvent b over 7 minutes at a flow rate of 0.2 mL/min and an injection volume of 5 μL with the oven temperature of 40°C. Analyte detection was achieved using a Shimadzu LC8030 triple quadrupole operating in ESI+ mode.

### Statistical Analysis

Statistical analysis was carried out in GraphPad Prism version 10.2.2 (GraphPad Software, La Jolla, CA). Comparison of Kaplan-Meier curves from the efficacy studies were carried out using the Log-rank (Mantel-Cox) test. For semi-quantitative analysis, the mean area between treatment groups was done using one-way ANOVA and Tukey’s post hoc test with a p value of less than 0.05 used to assess significance. For comparing mean areas between male and female pups within the same treatment group, an unpaired t-test was used with a p value of less than 0.05 used to assess significance. A t-test with Welch’s correction was done when comparing average DMA levels in the plasma between the 2 formulations with a p value of less than 0.05 used to determine significance. Additionally, an F test was done to compare variance between formulations with a p value of less than 0.05 used to determine significance.

## Results

### Delay of PTB with ip DMA vs. vaginal SNEDDS DMA administration

DMA administered ip consistently delayed LPS-induced PTB for 24 hours (Table 1). While 4 out of 8 mice in the control group delivered within 13 hours of LPS induction, none of the 10 mice treated with ip DMA delivered following ip LPS administration for the 24-hour experimental period (Figure 1, A). Pups that were delivered in the control group were expelled between 14 and 17 hours after the LPS induction (Figure 1, B). All the LPS-induced mice treated with the blank vaginal nanoformulation (n=6) similarly went into labor approximately 13-17 hours after treatment with ip LPS, while only 2 of the 8 LPS-induced mice given SNEDDS DMA went into labor. The 2 mice that delivered prematurely expelled their pups at 15-18 hours following LPS treatment (Figure 1, C), with SNEDDS DMA allowing all the pups to be retained in the uterine horns longer than the controls, including those that were delivered (Figure 1, D

**Figure 1.**
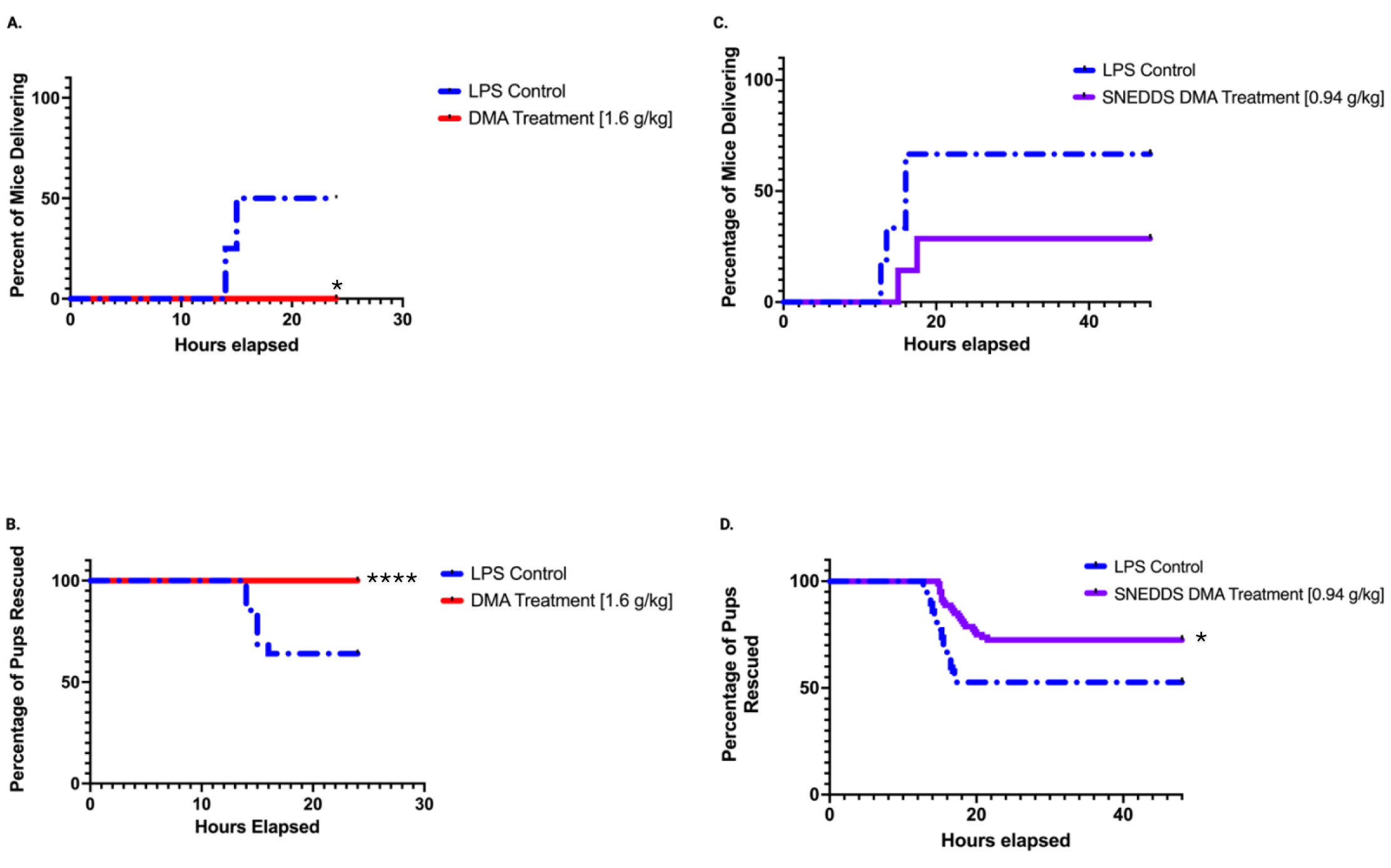
Delay and Prevention of LPS-Induced Preterm Birth with Intraperitoneal and SNEDDS DMA. The Kaplan-Meier Survival Curves looking at the percentage of mice delivering 24 hours after intraperitoneal LPS administration and treatment with IP DMA **(a)** and the percentage of pups that were retained in the inflamed uterus of pregnant CD-1 mice over a period of 24 hours following treatment with IP DMA **(b)**. The Kaplan-Meier Survival Curve looking at the percentage of mice delivering 48 following dosing with intraperitoneal LPS administration and treatment with SNEDDS DMA or a sham nanoformulation **(c)** and the percentage of pups that were retained in the inflamed uterus of pregnant CD-1 mice over a period of 48 hours after treatment **(d).**

**Table 1:** SNEDDS DMA Has Comparable Efficacy to Intraperitoneal DMA to Reduce the Incidence of Preterm Birth. Intraperitoneal DMA (IP DMA) was efficacious in preventing preterm birth in an inflammation-induced mouse model. Mice treated with i.p. DMA did not spontaneously abort any of their pups 24 hours following LPS administration. The DMA-loaded vaginal nanoformulation (SNEDDS DMA) was shown to delay and prevent LPS-induced preterm birth and retaining pups in utero at a lower dose.

|  | IP PBS +<br>LPS (1.6<br>mg/kg) | IP DMA (1.6<br>g/kg) + LPS<br>(1.6 mg/kg) | Blank<br>Nanoformulation +<br>LPS (1.6 mg/kg) | SNEDDS DMA (0.94<br>g/kg) + LPS (1.6<br>mg/kg) |
| --- | --- | --- | --- | --- |
| Number of Mice | 8 | 10 | 6 | 8 |
| Number of Mice<br>Delivering | 4 | 0*p<0.05 | 4 | 2 |
| Total Number of<br>Pups | 90 | 113 | 57 | 80 |
| Number of Pups<br>Delivered<br>Prematurely | 32 | 0*p<0.0001 | 27 | 22**p<0.05 |
| Number of Pups<br>Retained/Rescued<br>from Preterm Birth | 58 | 113 | 30 | 58 |

### Effects of ip DMA vs. vaginal SNEDDS DMA on placental and yolk sac histomorphology

At gestational day 17, the mouse yolk sac is characterized by elongated folds lined by cuboidal epithelial cells^38^ (endodermal cells). In mice treated with ip DMA (Figure 2, B), the structure of the cuboidal epithelial cells on the outermost layer of the yolk sac has been altered and the core of connective tissue underneath the endodermal cells is exposed. The blood islands in the mesoderm layer are now visible in ip DMA treated mice due to the endodermal cell layer being stripped away. These changes are attenuated in the yolk sacs of mice treated with vaginal SNEDDS DMA (Figure 2, D).

**Figure 2.**
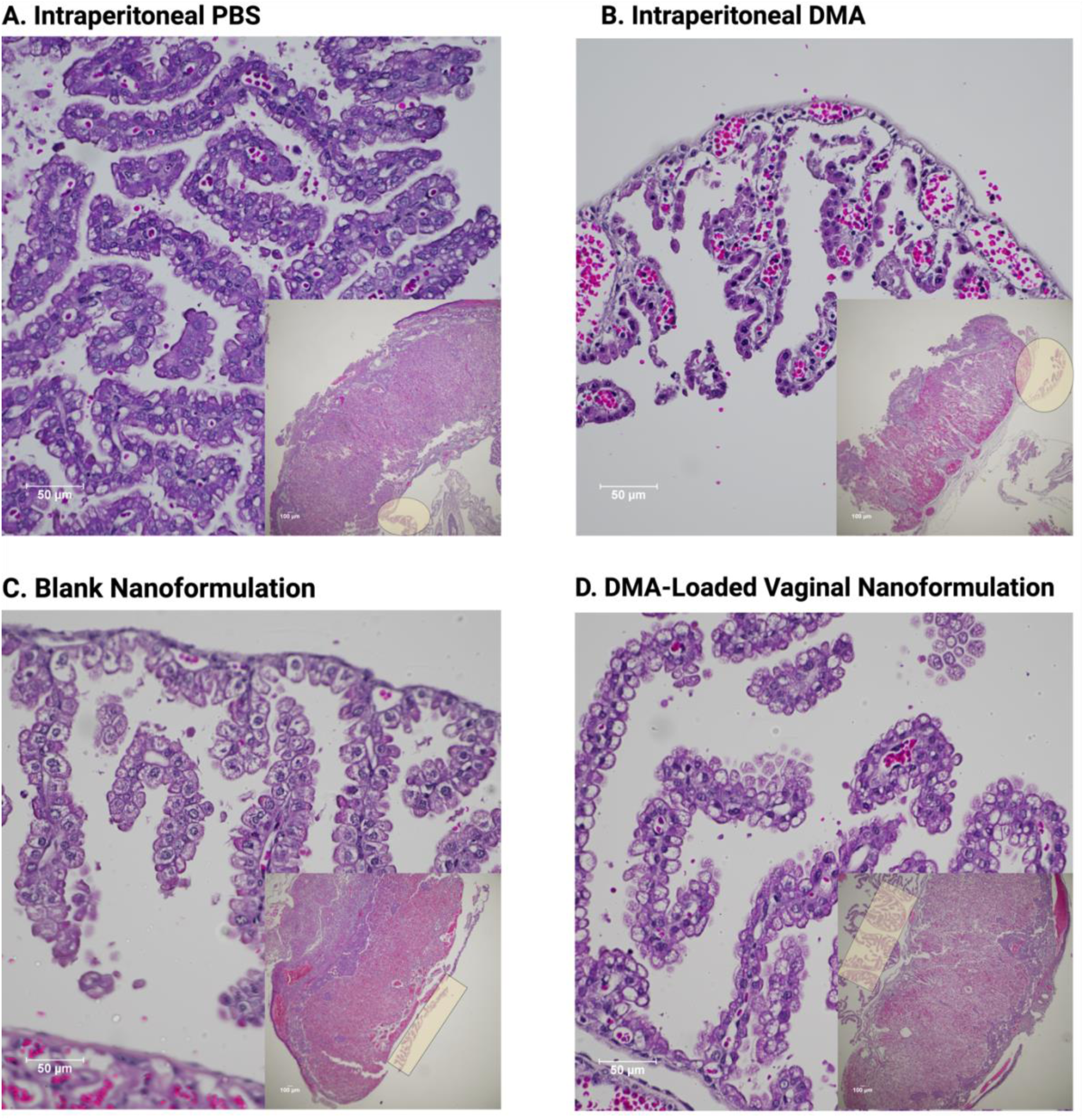
H&E Staining of the Yolk Sac following DMA Administration. Yolk Sacs were harvested from pregnant CD-1 mice on gestation day 17 and stained with H&E. Intraperitoneal administration of DMA compromises the architecture of the yolk sac, have deleterious effects not only on the superficial endodermal cells of the yolk sac but also on the thin region of loose connective tissue and blood vessels of the yolk sac mesoderm **[Panel B].** While there is some damage to the yolk sac architecture following administration of the DMA-Loaded Vaginal nanoformulation as seen in **[Panel D],** it is not to the same extent as intraperitoneal DMA (magnification 400x, scale bar 50 μM, n=6-8/group). **Insets:** Low magnification (40x, scale bar 100 μM) showing location of the yolk sac (highlighted) relative to the placenta it was attached to.

In the third trimester of mouse gestation, the labyrinth zone of the definitive placenta resembles highly branched cords surrounded by vascular channels, becoming more porous as parturition approaches^34,40,41^. Within the labyrinth, the biggest difference amongst the 4 treatment groups was the number of erythrocytes circulating in the vascular channels. Mice treated with ip DMA showed the biggest spike in volume of erythrocytes circulating in the vascular channels (Figure 3, B). Mice treated with vaginal SNEDDS DMA also exhibited labyrinth congestion but not to the same extent as ip DMA-treated mice. Based on semi-quantitative analysis with ImageJ/Fiji (Figure 3, I), the mean area of erythrocytes occupying the placental labyrinth in ip DMA treated mice significantly differed from mice treated with ip PBS (36536.9 μM^2^ vs. 22015.9 μM^2^; P<0.0001), the sham nanoformulation (36536.9 μM^2^ vs. 25570.2 μM^2^; P=0.0002), and SNEDDS DMA (36536.9 μM^2^ vs. 27864.3 μM^2^; P=0.0020).

**Figure 3.**
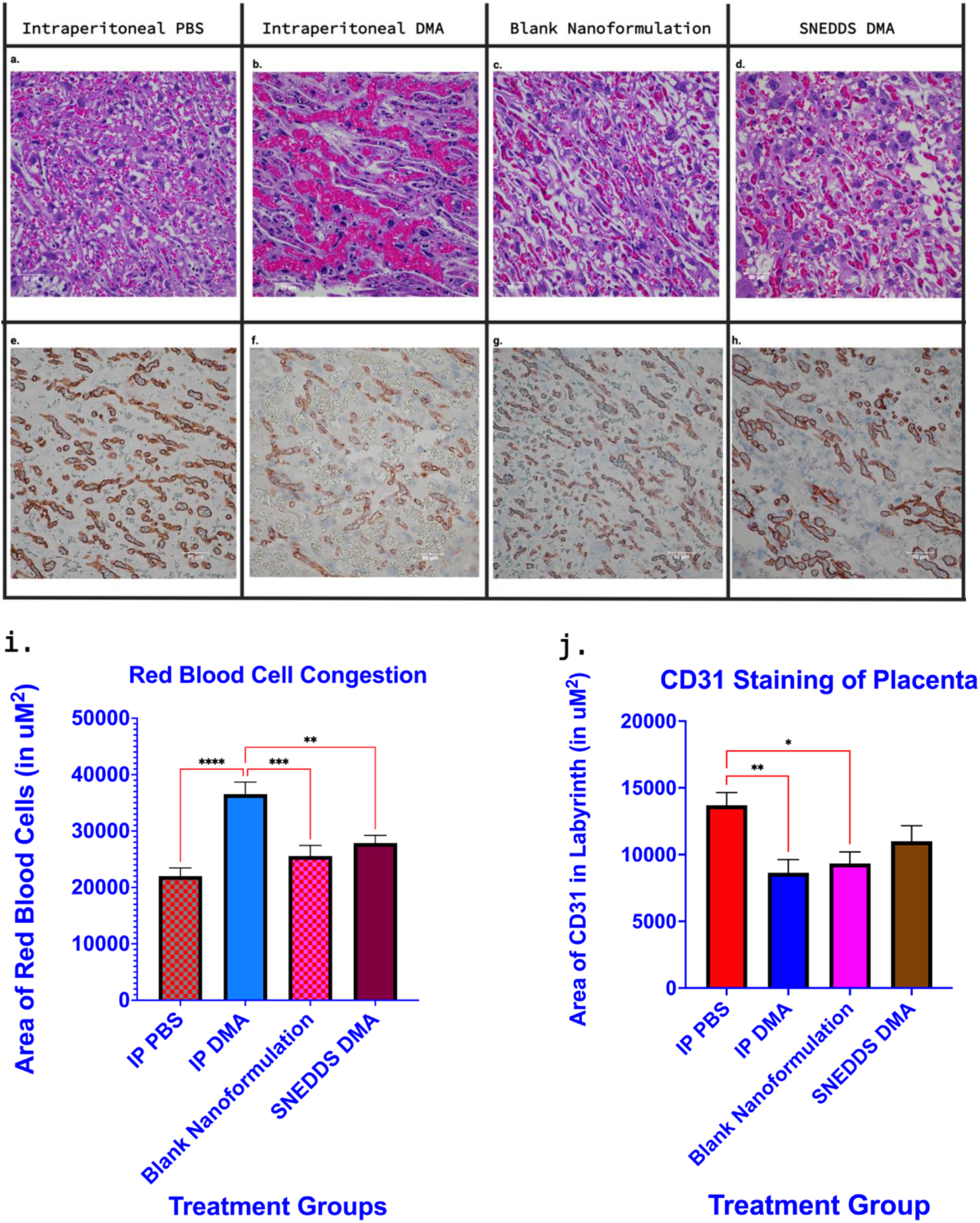
The effect of intraperitoneal and vaginal dimethylacetamide on the mouse placental labyrinth. Placentas were collected from CD1 dams on E17 after being treated with 3 doses of either i.p. DMA (1.6 g/kg), a vaginal nanoformulation of DMA (0.94 g/kg) or their respective negative controls, as indicated (n=21-28/group). Representative H&E-stained sections of the placental labyrinth from the four groups are shown in **panels A-D**. Systemic administration of DMA (i.p.) and SNEDDS DMA results in increased blood flow/congestion in the placental labyrinth, although to a lesser extent with SNEDDS DMA. Representative sections of the placental labyrinth stained with CD31 (brown) and counterstained with hematoxylin are shown in **panels E-H**. IP DMA-treated mice have decreased CD31 expression in the endothelial cells that make up the fetal blood vessels in the placental labyrinth. Semi-quantitative analysis (Mean and SEM in μM^2^) of villous congestion and CD31 content with ImageJ/Fiji is shown in **Panels I and J**, respectively. (Panels A-H Region of Interest magnification 400x, scale bar 50 μM).

Platelet/endothelial cell adhesion molecule (CD31) is known to be expressed at the junction between endothelial cells^41,42^. Between the maternal and fetal vascular channels in the labyrinth, only the fetal vasculature is lined by an endothelium^43^, therefore, this presents an opportunity to exclusively stain/visualize fetal endothelial cells in the labyrinth with CD31 antibody. Mice treated with ip DMA had lower CD31 expression compared to control mice treated with ip PBS (Figure 3, E-H). Quantitative analysis with ImageJ/Fiji (Figure 3, J) revealed the average area of CD31 staining in ip DMA-treated mice significantly differed from ip PBS-treated mice (≍ 8638 μM^2^ vs. ≍ 13700 μM^2^; P=0.034) while SNEDDS DMA exhibited no significant difference from the ip PBS group (≍ 11000 μM^2^ vs. ≍ 13700 μM^2^; P=0.2316).

After quantifying villous red blood cell congestion and CD31 staining between treatment groups, to address the variability within the same treatment group(s), another study was done to see if the severity of the 2 quantified adverse effects in the placenta, (red blood cell congestion and CD31 staining) vary with sex, that is, if it makes a difference if the placenta was attached to a male baby pup or a female baby pup. Red blood cell congestion and CD31 content was assessed between placentas attached to both sexes with individual and average values tabulated (Supplemental Figure 1). No significant differences were found between placentas attached to male pups and placentas attached to female pups.

### Systemic DMA concentration curves with ip DMA vs. vaginal SNEDDS DMA administration

DMA levels were assayed in plasma using quantitative high-performance liquid chromatography coupled to triple quadrupole mass spectrometry (LC/MS-MS) in MRM mode against standard curves. Both formulations had similar absorption profiles in that it takes approximately 60 minutes for DMA levels to reach their maximum plasma concentrations before beginning to decline (Figure 4, A). The average DMA concentrations were reduced by more than 50% in the systemic circulation when given SNEDDS DMA as opposed to ip DMA (Table 2) at 15 minutes following administration (5.47 mmol/L vs. 12.36 mmol/L; P=0.03) (Figure 4, B), 60 minutes following administration (5.88 mmol/L vs. 13.10 mmol/L; P =0.0004) (Figure 4, C), and 120 minutes following administration (4.95 mmol/L vs. 11.15 mmol/L; P=0.0005) (Figure 4, B). When comparing the variance associated with each route of administration using an F test, there was a significant difference in the concentration ranges between SNEDDS DMA and ip DMA at 15 minutes after administration (3.52 mmol/L – 9.82 mmol/L vs. 3.15 mmol/L – 23.59 mmol/L; P=0.0009), and 120 minutes after administration (4.04 mmol/L – 6.18 mmol/L vs. 7.62 mmol/L – 16.92).

**Figure 4.**
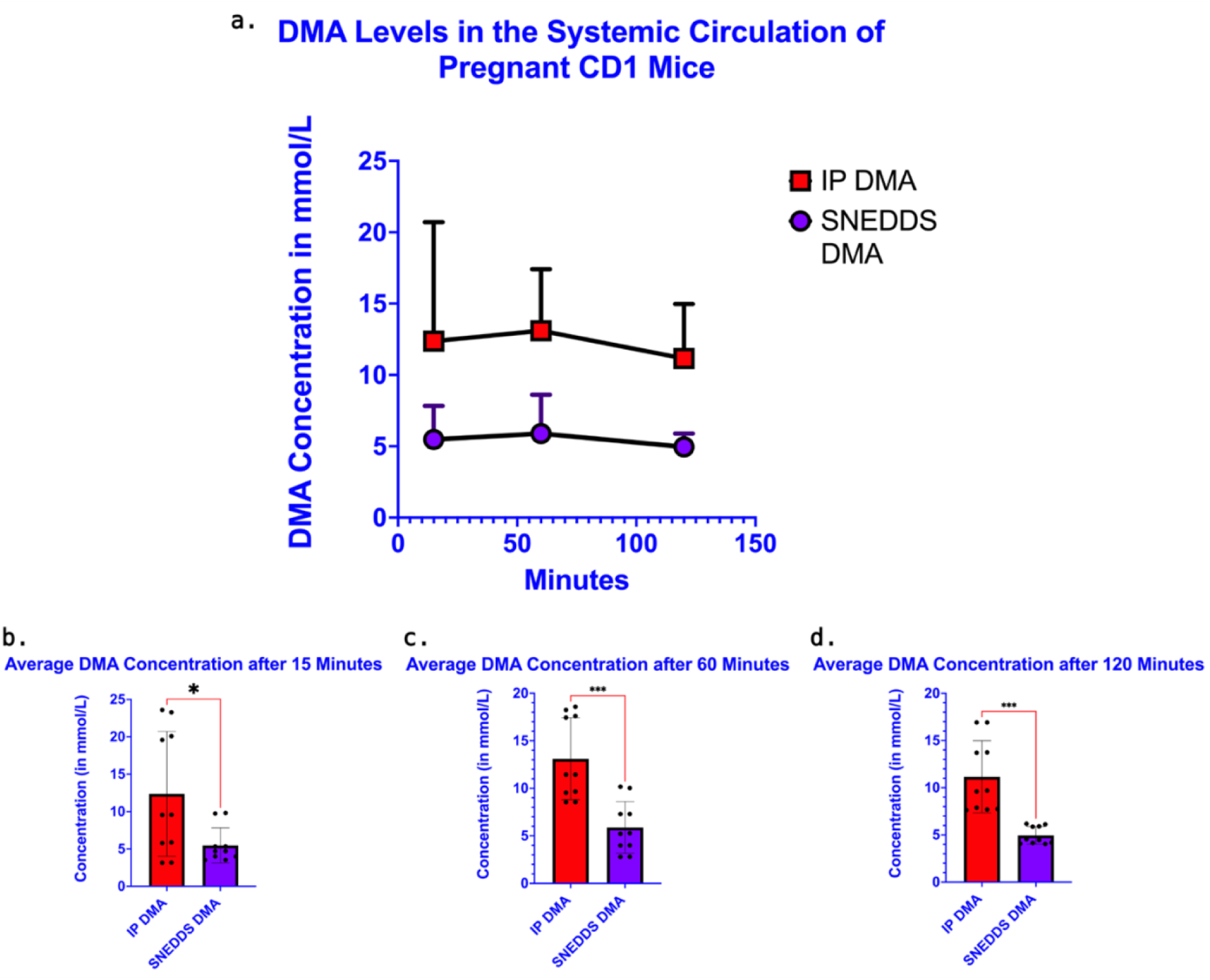
SNEDDS Formulation Reduces DMA Levels in the Systemic Circulation of Pregnant Mice. The concentration of DMA in plasma was quantified in E15 mice given a 1.56 g/kg intraperitoneal dose of DMA and E15 mice given a 1.56 g/kg vaginal dose of SNEDDS DMA at 3 different time points: 15, 60 & 120 minutes after administration **(a)**. At each time point, average concentrations in the SNEDDS treatment group were less than half the average concentration in the IP DMA treatment group. A student’s t-test shows SNEDDS DMA leads to a statistically significant reduction of DMA in the systemic circulation at 15 minutes after DMA administration **(b)**, 60 minutes after DMA administration **(c)**, and 120 minutes after DMA administration **(d).** (n=5/treatment group & time point, mean and SD in mmol/L shown)

**Table 2:**
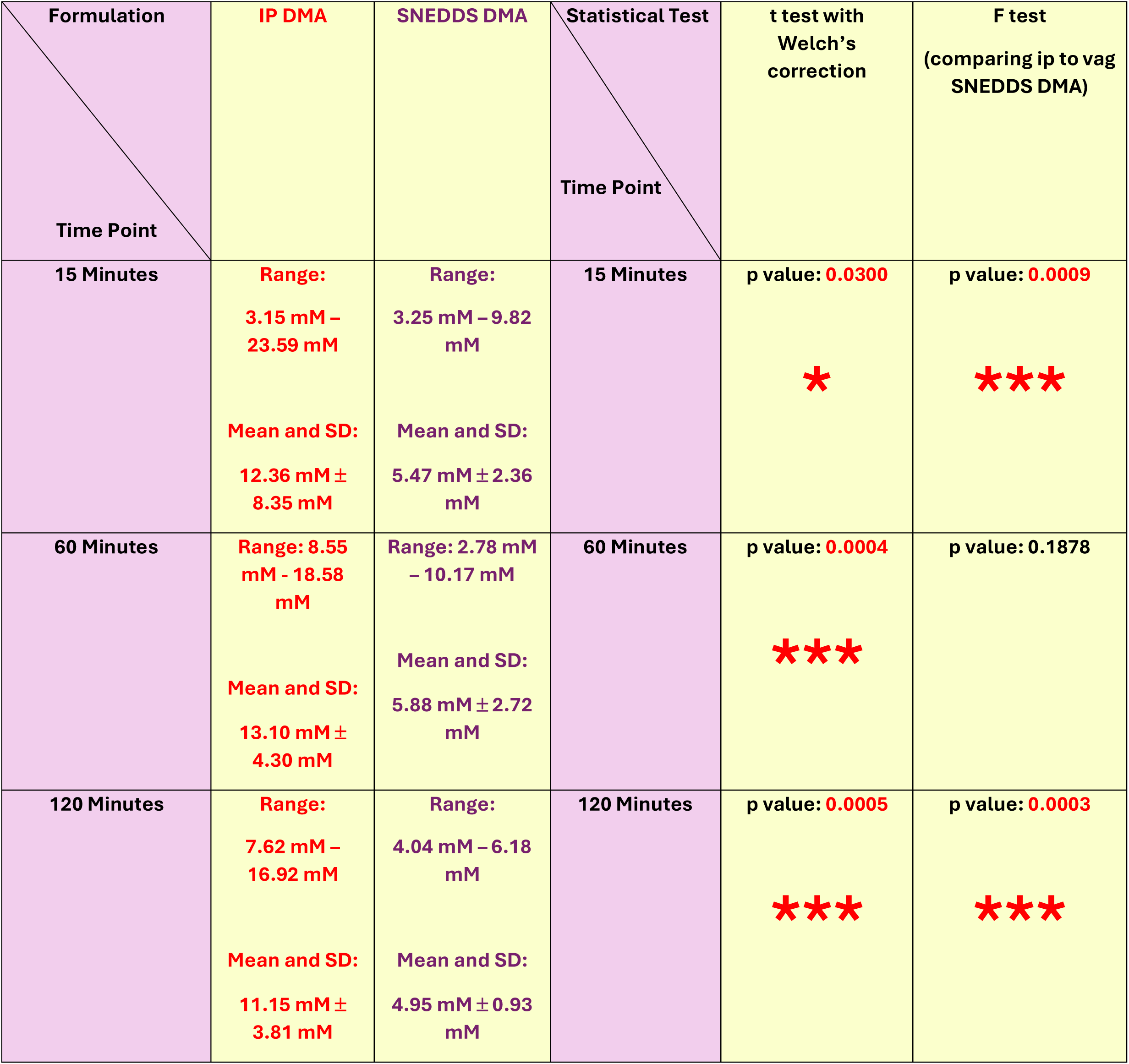
Statistical Analysis of DMA Concentrations in the Systemic Circulation via 2 Different Routes of Administration. A student’s t-test at each time point revealed the staggering difference in the average concentrations of DMA between both formulations with resulting p value increasing as time went on. The F-test also showed a significant difference in the large variability associated with intraperitoneal administration of DMA at the moment immediately following treatment and 120 minutes after treatment.

## Discussion

This side-by-side comparison of two different formulations of DMA given via two different routes of administration in a mouse PTB model demonstrates that (i) vaginal SNEDDS DMA has comparable efficacy to ip DMA at a lower dose (0.94 g/kg vs. 1.6 g/kg); (ii) effects of ip DMA on yolk sac morphology are attenuated with vaginal SNEDDS DMA administration; (iii) labyrinth congestion in the placenta is significantly decreased with vaginal SNEDDS DMA compared to ip DMA and loss of CD31 staining is alleviated with vaginal SNEDDS DMA; (iv) when given at the same dose as ip DMA, vaginal SNEDDS DMA reduces systemic levels of DMA by more than half; (v) ip DMA administration results in greater variation in systemic DMA concentration compared to vaginal SNEDDS DMA; (vi) sex does not play a role in the severity of ip DMA’s effects in the placenta.

Our lab’s first study testing DMA’s efficacy showed that at a dose of 1.56 g/kg (ip) prevents LPS-induced preterm birth altogether in C57BL/6 mice^32^. Similar results were obtained in this study with CD-1 mice, a strain with more genetic variability. Although other vaginal nanoformulations have been studied by our group and others^15,16^, a side-by-side comparison of efficacy, effects on placental and yolk sac histomorphology, and pharmacokinetics between an ip formulation and a vaginal SNEEDS has never been previously reported.

Regarding the efficacy of SNEDDS DMA, although 2 of the 8 mice delivered, this endpoint should not be interpreted as a sign of failure. In the 2 mice that delivered, arrival of the conceptus was delayed by up to 9 hours compared to controls, which roughly translates to almost 5 days in human gestation. This delay is significant as retaining the developing fetus in the uterus could allow for the proper development of tissues/organs and decrease the likelihood of neonatal morbidity and mortality. While both formulations caused labyrinth congestion, we believe this is not a reason to discard DMA as a potential treatment for preterm birth. The histology shows that although there is an increased number of circulating erythrocytes in the vascular channels, there is no hemorrhaging/rupture of blood vessels in the labyrinth. If there was damage to blood vessels, it would disqualify DMA as a viable therapeutic candidate for preterm birth. DMA causing a decrease in CD31 content in the labyrinth suggests a possible role in interfering with the development of fetal blood vessels. However, it is unknown if this is due to an insult to the placenta or secondary to an insult to another organ/tissue.

The decrease in systemic levels of DMA in mice treated with the vaginal SNEDDS formulation compared to ip DMA is due to the increased surface area and vascular network associated with vaginal administration^9,19^, allowing for enhanced delivery to the more pertinent tissues of the reproductive tract such as the uterus and cervix^10^. The uterine first-pass effect dictates that a significant portion of DMA, when given vaginally, is confined to the reproductive tract and minimizes the amount of DMA that makes it to the systemic circulation^45^. The former has not been proven yet, but the latter statement has been demonstrated in our study. In the context of treating preterm birth, elevated DMA levels in the systemic circulation via ip administration might pose a problem in that the tissues/organs of the mother which play a role in the development of the fetus, such as the kidneys, the liver, and the lungs are vulnerable^46–48^. This presents the possibility that adverse effects seen in the placenta and extra-embryonic tissues like the yolk sac following ip administration might be secondary to damage outside the reproductive tract, disrupting the crosstalk between the tissues of the mother and the developing tissues of the fetus, highlighting the potential advantage of vaginal administration over other routes of administration.

The results reported here have a very important implication for the PTB drug development field. The finding that a vaginal nanoformulation of a compound that prevents PTB *in vivo* has similar efficacy compared to its systemic formulation but attenuates changes in morphology at the fetal maternal interface suggests that drug therapies for PTB previously disqualified for toxicity issues can be revisited.

Our DMA vaginal SNEDDS formulation 1) takes advantage of the uterine first pass effect because of the vaginal route of administration of DMA and 2) decreases systemic DMA concentration by loading the DMA cargo into a nanoparticle. A strength of the study is the side-by-side comparison of the efficacy, effects on placental and yolk sac morphology and pharmacokinetics of the two DMA formulations. One minor weakness is that the pharmacokinetic study could be extended for a more precise estimate of DMA’s half-life^49–51^. Also, this study is limited to testing the maternal systemic circulation for DMA and does not extend the pharmacokinetic analysis to the fetal or placental compartments. Finally, while beyond the scope of this proof of principle study which tests the formulations for their effect on length of gestation, a comparison of the two approaches on neonatal outcomes in liveborn pups will be needed to determine which formulation is more suited for clinical application.

Our study shows that SNEDDS DMA is a viable and even favorable alternative to systemic DMA when considering potential treatments for preterm birth. In late gestation, not only does it have comparable efficacy to systemic DMA, but it can curtail histomorphological effects seen in the reproductive tract when DMA is given ip. Overall, this study adds to the growing body of literature that suggests vaginal nanoformulations are the breakthrough technology in the field of reproductive medicine that will be able to tackle the urgent issue of preterm birth.

## Acknowlegments

The authors would like to thank Fangping Chen, the managing director of the histotechnology facility at the Wistar Institute located in Philadelphia, Pennsylvania for her expertise and assistance for the histology and immunohistochemical staining of tissues/specimens.

## Author Contributions

S.E.R., C.M., S.M.Y., and K.P. designed and supervised the study. A.M. and T.A. performed the experiments and analyzed the data.

## Declaration of Interests

SER has been issued a patent titled “Administration of N,N-Dimethylacetamide for the Treatment of Preterm Birth” (Patent Number US 11,717,499 B2).

## Funding

This work was supported by the National Institutes of Health (Grant 1R16GM145586).

## Notes

### Competing Interest Statement

The authors have declared no competing interest.

## References

1. Harris, E. (2023). FDA Revokes Approval for Preterm Birth Drug Makena. JAMA, 329(17), 1444– 1444. 10.1001/jama.2023.6257

2. Khandre, V., Potdar, J., & Keerti, A. (2022). Preterm Birth: An Overview. Cureus, 14(12), e33006. 10.7759/cureus.33006

3. Manuck T. A. (2017). Racial and ethnic differences in preterm birth: A complex, multifactorial problem. Seminars in perinatology, 41(8), 511–518. 10.1053/j.semperi.2017.08.010

4. Blackwell, S. C., Gyamfi-Bannerman, C., Biggio, J. R., Jr, Chauhan, S. P., Hughes, B. L., Louis, J. M., Manuck, T. A., Miller, H. S., Das, A. F., Saade, G. R., Nielsen, P., Baker, J., Yuzko, O. M., Reznichenko, G. I., Reznichenko, N. Y., Pekarev, O., Tatarova, N., Gudeman, J., Birch, R., Jozwiakowski, M. J., Duncan, M., Williams, L., Krop, J. (2020). 17-OHPC to Prevent Recurrent Preterm Birth in Singleton Gestations (PROLONG Study): A Multicenter, International, Randomized Double-Blind Trial. American journal of perinatology, 37(2), 127–136. 10.1055/s-0039-3400227

5. Nelson, D. B., Herrera, C. L., McIntire, D. D., & Cunningham, F. G. (2024). The end is where we start from: Withdrawal of 17-alpha hydroxyprogesterone caproate to prevent recurrent preterm birth. American Journal of Obstetrics & Gynecology, 230(1), 1–9. 10.1016/j.ajog.2023.08.031

6. Alexander, N. J., Baker, E., Kaptein, M., Karck, U., Miller, L., & Zampaglione, E. (2004). Why consider vaginal drug administration? Fertility and Sterility, 82(1), 1–12. 10.1016/j.fertnstert.2004.01.025

7. Cicinelli, E., Di Naro, E., De Ziegler, D., Matteo, M., Morgese, S., Galantino, P., Brioschi, P. A., & Schonauer, A. (2003). Placement of the vaginal 17beta-estradiol tablets in the inner or outer one third of the vagina affects the preferential delivery of 17beta-estradiol toward the uterus or periurethral areas, thereby modifying efficacy and endometrial safety. American journal of obstetrics and gynecology, 189(1), 55–58. 10.1067/mob.2003.341

8. Zierden, H. C., Shapiro, R. L., DeLong, K., Carter, D. M., & Ensign, L. M. (2021). Next generation strategies for preventing preterm birth. Advanced drug delivery reviews, 174, 190–209. 10.1016/j.addr.2021.04.021

9. Mir, A., Vartak, R. V., Patel, K., Yellon, S. M., & Reznik, S. E. (2022). Vaginal Nanoformulations for the Management of Preterm Birth. Pharmaceutics, 14(10), 2019. 10.3390/pharmaceutics14102019

10. Bulletti, C., de Ziegler, D., Flamigni, C., Giacomucci, E., Polli, V., Bolelli, G., & Franceschetti, F. (1997). Targeted drug delivery in gynaecology: the first uterine pass effect. Human reproduction (Oxford, England), 12(5), 1073–1079. 10.1093/humrep/12.5.1073

11. Hussain, A., & Ahsan, F. (2005). The vagina as a route for systemic drug delivery. Journal of controlled release: official journal of the Controlled Release Society, 103(2), 301–313. 10.1016/j.jconrel.2004.11.034

12. Ensign, L. M., Tang, B. C., Wang, Y. Y., Tse, T. A., Hoen, T., Cone, R., & Hanes, J. (2012). Mucus-penetrating nanoparticles for vaginal drug delivery protect against herpes simplex virus. Science translational medicine, 4(138), 138ra79. 10.1126/scitranslmed.3003453

13. Fida, S., Jalil, A., Habib, R., Akhlaq, M., Mahmood, A., Minhas, M. U., Khan, K. U., & Nawaz, A. (2022). Development of mucus-penetrating iodine loaded self-emulsifying system for local vaginal delivery. PloS one, 17(3), e0266296. 10.1371/journal.pone.0266296

14. Pedersen, C., Slepenkin, A., Andersson, S. B., Fagerberg, J. H., Bergström, C. A., & Peterson, E. M. (2014). Formulation of the microbicide INP0341 for in vivo protection against a vaginal challenge by Chlamydia trachomatis. PloS one, 9(10), e110918. 10.1371/journal.pone.0110918

15. Patki, M., Giusto, K., Gorasiya, S., Reznik, S. E., & Patel, K. (2019). 17-α Hydroxyprogesterone Nanoemulsifying Preconcentrate-Loaded Vaginal Tablet: A Novel Non-Invasive Approach for the Prevention of Preterm Birth. Pharmaceutics, 11(7), 335. 10.3390/pharmaceutics11070335

16. Giusto, K., Patki, M., Koya, J., Ashby, C. R., Jr, Munnangi, S., Patel, K., & Reznik, S. E. (2019). A vaginal nanoformulation of a SphK inhibitor attenuates lipopolysaccharide-induced preterm birth in mice. Nanomedicine (London, England), 14(21), 2835–2851. 10.2217/nnm-2019-0243

17. Lacroix, G., Gouyer, V., Gottrand, F., & Desseyn, J. L. (2020). The Cervicovaginal Mucus Barrier. International journal of molecular sciences, 21(21), 8266. 10.3390/ijms21218266

18. Zierden, H. C., DeLong, K., Zulfiqar, F., Ortiz, J. O., Laney, V., Bensouda, S., Hernández, N., Hoang, T. M., Lai, S. K., Hanes, J., Burke, A. E., & Ensign, L. M. (2023). Cervicovaginal mucus barrier properties during pregnancy are impacted by the vaginal microbiome. Frontiers in Cellular and Infection Microbiology, 13. 10.3389/fcimb.2023.1015625

19. Srikrishna, S., & Cardozo, L. (2013). The vagina as a route for drug delivery: a review. International urogynecology journal, 24(4), 537–543. 10.1007/s00192-012-2009-3

20. Silvia Rossi, C. M. C., Barbara Vigani, Giuseppina Sandri, Maria Cristina Bonferoni, & Ferrari, F. (2019). Recent advances in the mucus-interacting approach for vaginal drug delivery: From mucoadhesive to mucus-penetrating nanoparticles. Expert Opinion on Drug Delivery, 16(8), 777– 781. 10.1080/17425247.2019.1645117

21. Zhang, Y., Li, S., Loch, K., Duncan, G. A., Kaler, L., Pangeni, R., Peng, W., Wang, S., Gong, X., & Xu, Q. (2023). PH-Responsive Mucus-Penetrating Nanoparticles for Enhanced Cellular Internalization by Local Administration in Vaginal Tissue. ACS Macro Letters, 12(4), 446–453. 10.1021/acsmacrolett.2c00639

22. Taylor, J., Sharp, A., Rannard, S. P., Arrowsmith, S., & McDonald, T. O. (2023). Nanomedicine strategies to improve therapeutic agents for the prevention and treatment of preterm birth and future directions. Nanoscale advances, 5(7), 1870–1889. 10.1039/d2na00834c

23. Smoleński, M., Karolewicz, B., Gołkowska, A. M., Nartowski, K. P., & Małolepsza-Jarmołowska, K. (2021). Emulsion-Based Multicompartment Vaginal Drug Carriers: From Nanoemulsions to Nanoemulgels. International journal of molecular sciences, 22(12), 6455. 10.3390/ijms22126455

24. Jahangirian, H., Kalantari, K., Izadiyan, Z., Rafiee-Moghaddam, R., Shameli, K., & Webster, T. J. (2019). A review of small molecules and drug delivery applications using gold and iron nanoparticles. International journal of nanomedicine, 14, 1633–1657. 10.2147/IJN.S184723

25. Jia, Y., Jiang, Y., He, Y., Zhang, W., Zou, J., Magar, K. T., Boucetta, H., Teng, C., & He, W. (2023). Approved Nanomedicine against Diseases. Pharmaceutics, 15(3), 774. 10.3390/pharmaceutics15030774

26. Patel, V. R., & Agrawal, Y. K. (2011). Nanosuspension: An approach to enhance solubility of drugs. Journal of advanced pharmaceutical technology & research, 2(2), 81–87. 10.4103/2231-4040.82950

27. Kyathanahalli, C., Snedden, M., & Hirsch, E. (2023). Is human labor at term an inflammatory condition? Biology of reproduction, 108(1), 23–40. 10.1093/biolre/ioac182

28. Norman, J. E., Bollapragada, S., Yuan, M., & Nelson, S. M. (2007). Inflammatory pathways in the mechanism of parturition. BMC pregnancy and childbirth, 7 Suppl 1(Suppl 1), S7. 10.1186/1471-2393-7-S1-S7

29. Cooley, A., Madhukaran, S., Stroebele, E., Colon Caraballo, M., Wang, L., Akgul, Y., Hon, G. C., & Mahendroo, M. (2023). Dynamic states of cervical epithelia during pregnancy and epithelial barrier disruption. iScience, 26(2), 105953. 10.1016/j.isci.2023.105953

30. Edey, L. F., Georgiou, H., O’Dea, K. P., Mesiano, S., Herbert, B. R., Lei, K., Hua, R., Markovic, D., Waddington, S. N., MacIntyre, D., Bennett, P., Takata, M., & Johnson, M. R. (2017). Progesterone, the maternal immune system and the onset of parturition in the mouse†. Biology of Reproduction, 98(3), 376–395. 10.1093/biolre/iox146

31. Pekson, R., Poltoratsky, V., Gorasiya, S., Sundaram, S., Ashby, C. R., Vancurova, I., & Reznik, S. E. (2016). N,N-Dimethylacetamide Significantly Attenuates LPS- and TNFα-Induced Proinflammatory Responses Via Inhibition of the Nuclear Factor Kappa B Pathway. Molecular medicine (Cambridge, Mass.), 22, 747–758. 10.2119/molmed.2016.00017

32. Sundaram, S., Ashby, C. R., Jr, Pekson, R., Sampat, V., Sitapara, R., Mantell, L., Chen, C. H., Yen, H., Abhichandani, K., Munnangi, S., Khadtare, N., Stephani, R. A., & Reznik, S. E. (2013). N,N-dimethylacetamide regulates the proinflammatory response associated with endotoxin and prevents preterm birth. The American journal of pathology, 183(2), 422–430. 10.1016/j.ajpath.2013.05.006

33. Bolon, B. (2014). Protocols for Placental Histology. In Croy, B. A., Yamada, A. T., DeMayo, F. J., & Adamson, S. L (Eds.) The Guide to Investigation of Mouse Pregnancy (1st ed., pp. 539). Elsevier/Academic Press.

34. Xu, B., Chen, X., Ding, Y., Chen, C., Liu, T., & Zhang, H. (2020). Abnormal angiogenesis of placenta in progranulin-deficient mice. Molecular medicine reports, 22(4), 3482–3492. 10.3892/mmr.2020.11438

35. Pikor LA, Enfield KS, Cameron H, Lam WL. DNA extraction from paraffin embedded material for genetic and epigenetic analyses. J Vis Exp. 2011 Mar 26;(49):2763. doi: 10.3791/2763. PMID: 21490570; PMCID: PMC3197328.

36. Heimann M, Roth DR, Ledieu D, Pfister R, Classen W. Sublingual and submandibular blood collection in mice: a comparison of effects on body weight, food consumption and tissue damage. Laboratory Animals. 2010;44(4):352–358. doi:10.1258/la.2010.010011

37. Rosser, S. P. A., McLachlan, A. J., Hempel, G., Chung, J., Shaw, P. J., Keogh, S. J., & Nath, C. E. (2023). Validation of a liquid chromatography-tandem mass spectrometry method for simultaneous quantification of N,N-dimethylacetamide and N-monomethylacetamide in pediatric plasma. Journal of separation science, 46(10), e2201003. 10.1002/jssc.202201003

38. Yamane T. (2018). Mouse Yolk Sac Hematopoiesis. Frontiers in cell and developmental biology, 6, 80. 10.3389/fcell.2018.00080

39. Hu, D., & Cross, J. C. (2010). Development and function of trophoblast giant cells in the rodent placenta. The International journal of developmental biology, 54(2-3), 341–354. 10.1387/ijdb.082768dh

40. Mu, J., & Adamson, S. L. (2006). Developmental changes in hemodynamics of uterine artery, utero- and umbilicoplacental, and vitelline circulations in mouse throughout gestation. American journal of physiology. Heart and circulatory physiology, 291(3), H1421–H1428. 10.1152/ajpheart.00031.2006

41. Vanchinathan, V., Mizramani, N., Kantipudi, R., Schwartz, E. J., & Sundram, U. N. (2015). The Vascular Marker CD31 Also Highlights Histiocytes and Histiocyte-Like Cells Within Cutaneous Tumors. American Journal of Clinical Pathology, 143(2), 177–185. 10.1309/AJCPRHM8CZH5EMFD

42. Cheung, K., Ma, L., Wang, G., Coe, D., Ferro, R., Falasca, M., Buckley, C. D., Mauro, C., & Marelli-Berg, F. M. (2015). CD31 signals confer immune privilege to the vascular endothelium. Proceedings of the National Academy of Sciences, 112(43), E5815–E5824. 10.1073/pnas.1509627112

43. Bolon, B. (2014). Pathology Analysis of the Placenta. In Croy, B. A., Yamada, A. T., DeMayo, F. J., & Adamson, S. L (Eds.) The Guide to Investigation of Mouse Pregnancy (1st ed., pp. 180). Elsevier/Academic Press.

44. Cicinelli, E., Ziegler, D. de, Morgese, S., Bulletti, C., Luisi, D., & Schonauer, L. M. (2004). “First uterine pass effect” is observed when estradiol is placed in the upper but not lower third of the vagina. Fertility and Sterility, 81(5), 1414–1416. 10.1016/j.fertnstert.2003.12.016

45. De Ziegler, D., Bulletti, C., De Monstier, B., & Jääskeläinen, A. S. (1997). The first uterine pass effect. Annals of the New York Academy of Sciences, 828, 291–299. 10.1111/j.1749-6632.1997.tb48550.x

46. Cabarcas-Barbosa, O., Capalbo, O., Ferrero-Fernández, A., & Musso, C. G. (2022). Kidney-placenta crosstalk in health and disease. Clinical kidney journal, 15(7), 1284–1289. 10.1093/ckj/sfac060

47. Lakhal-Littleton S. (2021). Advances in understanding the crosstalk between mother and fetus on iron utilization. Seminars in hematology, 58(3), 153–160. 10.1053/j.seminhematol.2021.06.003

48. Hansen, S. S. K., Krautz, R., Rago, D., Havelund, J., Stigliani, A., Færgeman, N. J., Prézelin, A., Rivière, J., Couturier-Tarrade, A., Akimov, V., Blagoev, B., Elfving, B., Neess, D., Vogel, U., Khodosevich, K., Hougaard, K. S., & Sandelin, A. (2024). Pulmonary maternal immune activation does not cross the placenta but leads to fetal metabolic adaptation. Nature Communications, 15(1), 4711. 10.1038/s41467-024-48492-x

49. Trame, M. N., Bartelink, I. H., Boos, J., Boelens, J. J., & Hempel, G. (2013). Population pharmacokinetics of dimethylacetamide in children during standard and once-daily IV busulfan administration. Cancer chemotherapy and pharmacology, 72(5), 1149–1155. 10.1007/s00280-013-2284-9

50. Hempel, G., Oechtering, D., Lanvers-Kaminsky, C., Klingebiel, T., Vormoor, J., Gruhn, B., & Boos, J. (2007). Cytotoxicity of Dimethylacetamide and Pharmacokinetics in Children Receiving Intravenous Busulfan. Journal of Clinical Oncology, 25(13), 1772–1778. 10.1200/JCO.2006.08.8807

51. Hundley, S. G., Lieder, P. H., Valentine, R., McCooey, K. T., & Kennedy, G. L., Jr (1994). Dimethylacetamide pharmacokinetics following inhalation exposures to rats and mice. Toxicology letters, 73(3), 213–225. 10.1016/0378-4274(94)90061-2

