## Supplemental Figures 1 and 2 for "Improving the Safety of *N,N*-Dimethylacetamide (DMA) as a Potential Treatment for Preterm Birth in a Pregnant Mouse Model Using a Vaginal Nanoformulation"

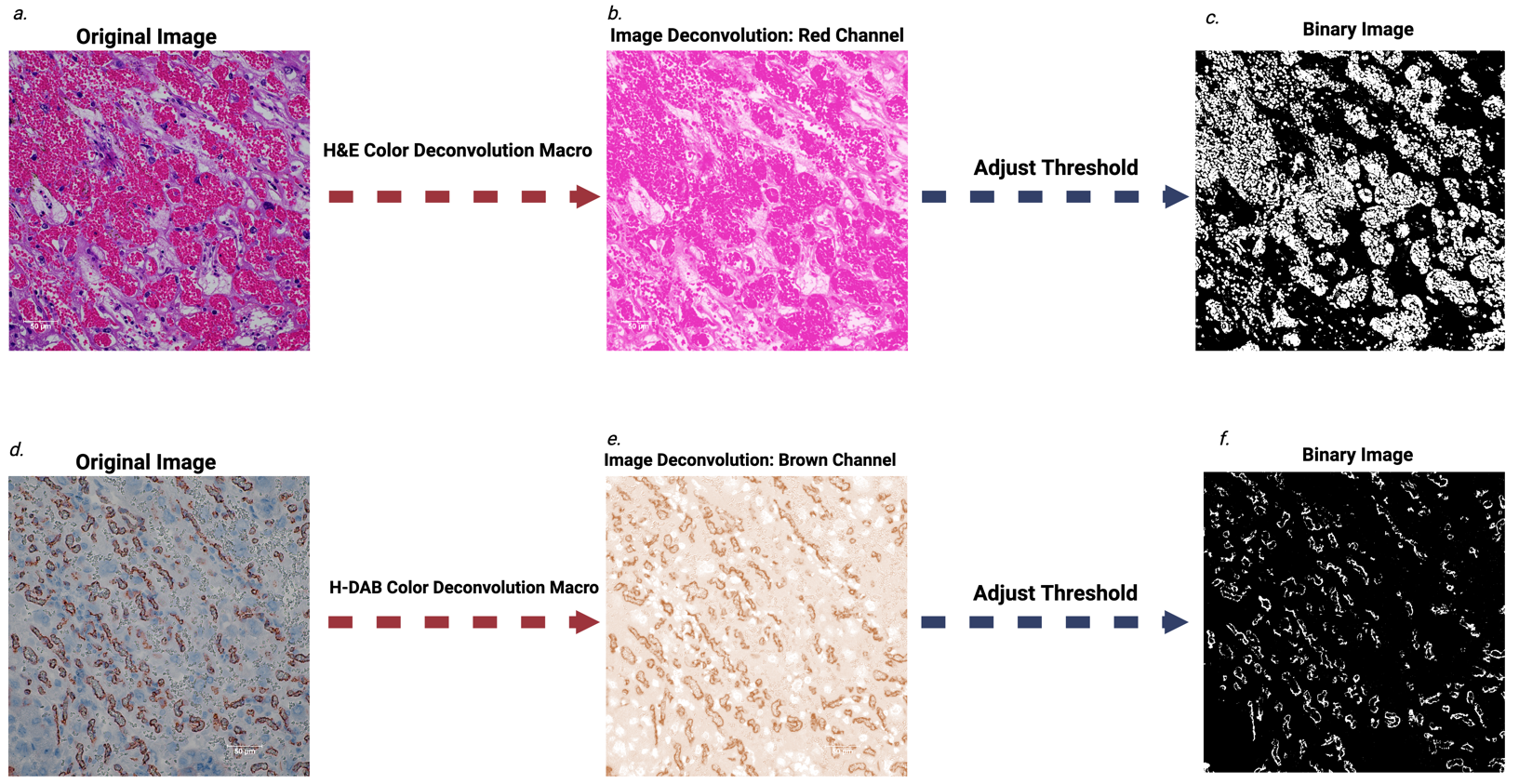

**Supplemental Figure 1: *Semi-quantitative histology workflow.*** *Sections were analyzed with a Nikon Eclipse Ts2R microscope and micrographs were captured with a Nikon Digital Sight 10 camera. Digitized images (400x magnification) were loaded into ImageJ/Fiji version 2.14.0 and underwent image deconvolution where they were separated into their respective channels and background subtraction was done by subtracting out pixels that did not fall within a range of color/pixel “intensities” to generate a binary image where the area of the pixels of interest can be measured. The “H&E Color Deconvolution Macro” was used for red blood cell area count as depicted in* ***panels A-C****. The “H-DAB Color Deconvolution Macro” was used for CD31 area count as depicted in* ***panels D-F.***

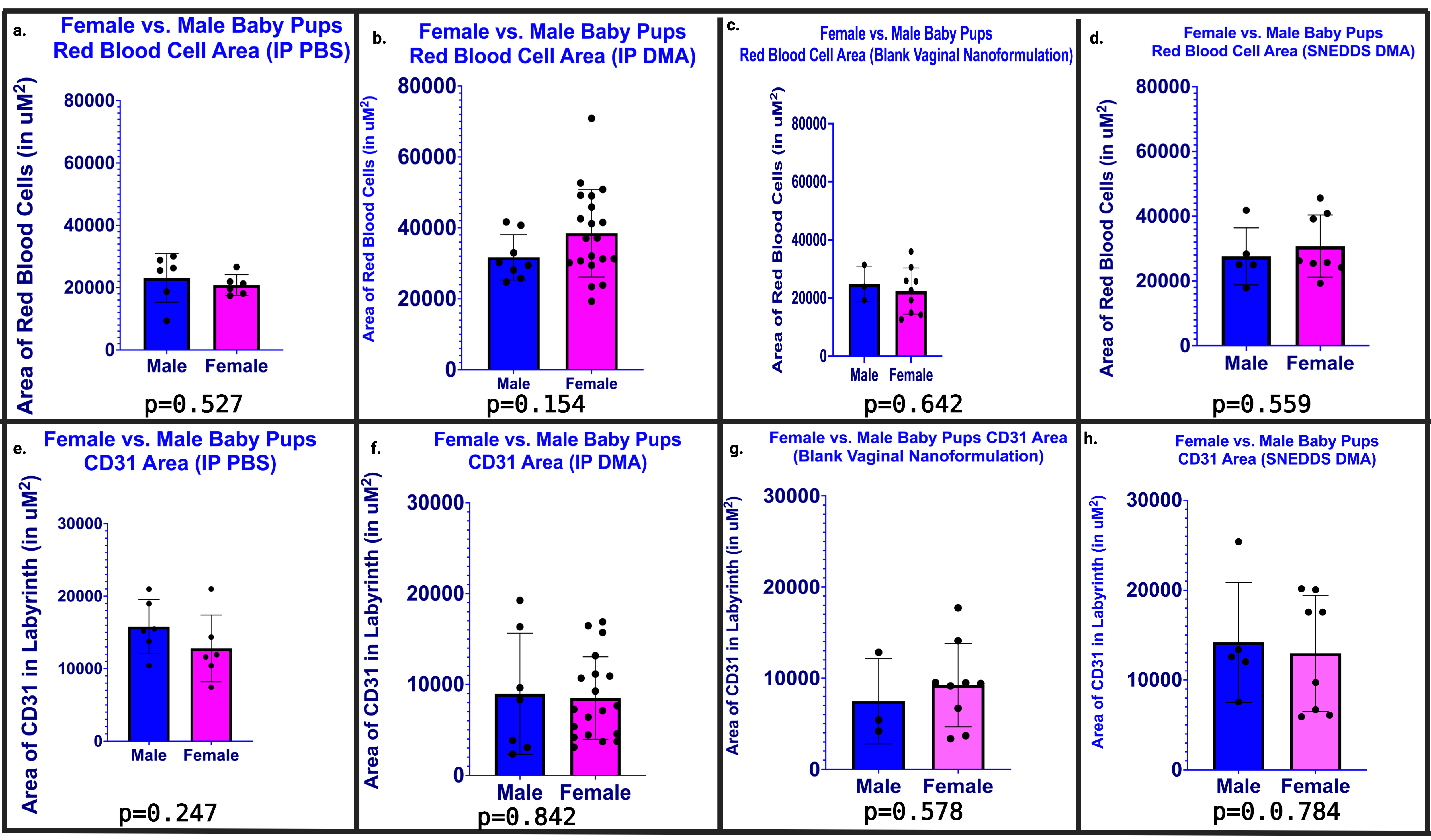

**Supplemental Figure 2: Sexual dimorphism does not play a role in DMA’s effect on placental histomorphology.** *Red blood cell areas were calculated in placental sections from each of the four groups, as indicated, and results for male and female palcentas were compared*  [***Panels A-D].*** *Similarly, CD31staining areas were calculated in placental sections from each of the four groups, as indicated, and results for male and female palcentas were compared* ***[Panels E-H].*** *(N=12-28/treatment group.)*
